# Maternal antibodies provide strain-specific protection against infection with the Lyme disease pathogen in a wild rodent

**DOI:** 10.1101/522789

**Authors:** Andrea Gomez-Chamorro, Vanina Heinrich, Anouk Sarr, Owen Roethlisberger, Dolores Genné, Maxime Jacquet, Maarten J. Voordouw

## Abstract

The vertebrate immune system can produce antibodies that protect the host against pathogens. Females can transmit antibodies to their offspring, which provide short-term protection against infection. The tick-borne bacterium *Borrelia afzelii* causes Lyme disease in Europe and consists of multiple strains that cycle between the tick vector (*Ixodes ricinus*) and vertebrate hosts such as the bank vole (*Myodes glareolus*). We used a controlled experiment to show that infected female bank voles transmit protective antibodies to their offspring that are specific for the strain of *B. afzelii*. To test the specificity of protection, the offspring were challenged with either the same strain to which the mothers had been exposed or a different strain. The maternal antibodies protected the offspring against the same strain, but not against the different strain. The offspring from the uninfected control mothers were equally susceptible to both strains. Our study shows that maternal antibodies provide strong but highly strain-specific protection against *B. afzelii* in an important rodent reservoir host. The transmission of maternal antibodies may have important consequences for the epidemiology of multiple-strain pathogens in nature.

**Author Summary:** Many pathogens that cause infectious disease consist of multiple strains. In vertebrate hosts, the immune system can generate antibodies that are highly specific for different pathogen strains. Mothers can transmit these antibodies to their offspring and thereby protect them from infectious disease. To date, few studies have investigated whether this transgenerational transfer of protective antibodies is important for pathogens that cycle in wild animal populations. The tick-borne spirochete bacterium *Borrelia afzelii* causes Lyme disease in Europe and cycles between *Ixodes* ticks and wild rodent hosts, such as the bank vole (*Myodes glareolus*). The purpose of our study was to test whether female bank voles infected with *B. afzelii* transmit antibodies to their offspring that protect them from an infected tick bite. Our study found that infected mothers do transmit antibodies, but the offspring were only protected against the strain of *B. afzelii* to which their mothers had been exposed and not to a different strain (i.e. protection was highly strain-specific). The broader implications of our study is that the transfer of protective antibodies between generations in the vertebrate host population could be important for organizing the community of pathogen strains that circulate in nature.

## Introduction

Parents transmit to their offspring more than just their genes [1]. Mothers transfer their environmental experience and phenotype to their offspring [2] and these maternal effects can influence offspring phenotype and fitness [3]. In vertebrate hosts, an important maternal effect is the transmission of antibodies from mothers to offspring [4]. Young vertebrates are susceptible to infectious diseases while their immune systems are developing [4]. Maternally transmitted antibodies can protect the offspring against pathogens until they can start to produce their own antibodies [4]. The transgenerational transmission of acquired immunity can have important consequences for the epidemiology of the pathogen [1]. For example, maternal antibodies were shown to influence the epidemiology of Puumala virus in wild bank vole populations [5]. Despite its potential importance in nature, the transgenerational transfer of acquired immunity has not received much attention in the epidemiology of zoonotic diseases [1,5].

Maternal antibodies may be particularly important for the epidemiology of pathogens that consist of multiple genetically distinct strains that circulate in the host population at the same time. These strains can be distinguished by the acquired immune system of the vertebrate host resulting in the development of strain-specific antibody responses. Theoretical models have shown that strain-specific antibodies play a critical role in shaping the epidemiology and population structure of pathogen strains [6,7,8,9]. Maternal antibodies are expected to be particularly important in systems where the vertebrate host is short-lived and has multiple generations within the transmission season of the pathogen. In such systems, a pathogen strain that is common in the maternal generation would be at a selective disadvantage in the offspring generation due to the maternal transmission of strain-specific antibodies against this common strain.

Tick-borne spirochete bacteria belonging to the *Borrelia burgdorferi* sensu lato (sl) genospecies complex are the etiological agents of Lyme borreliosis [10,11]. *B. burgdorferi* sl is a good model system for studying whether maternally transmitted antibodies can influence strain-specific infection success. The populations of *B. burgdorferi* sl consist of multiple strains that circulate between *Ixodes* ticks and vertebrate hosts such as rodents and birds [12,13,14,15,16]. Immature *Ixodes* ticks search for a blood meal from spring until early autumn, and the transmission of *B. burgdorferi* sl therefore coincides with the reproduction and population expansion of their vertebrate hosts [11,17]. There is no vertical transmission of *B. burgdorferi* sl in either the tick [18] or the vertebrate host [19,20]. In nature, vertebrate hosts develop a strong antibody response against *B. burgdorferi* sl [19,20], and infection studies have shown that this antibody response is strain-specific [21,22,23]. Previous field studies on a marine Lyme borreliosis system found that infected female seabirds transmit antibodies to their offspring [24,25]. However, to date no one has provided experimental evidence that maternal antibodies protect offspring against infection with *B. burgdorferi* sl and that this protection is strain-specific.

In this study, we used *Borrelia afzelii*, which is the most common cause of Lyme borreliosis in Europe [26], its tick vector *I. ricinus*, and the bank vole (*Myodes glareoulus*), which is an important reservoir host for both *B. afzelii* and *I. ricinus* [27,28,29]. The purpose of this study was to test (1) whether female bank voles that were experimentally infected with *B. afzelii* transmit maternal antibodies to their offspring, (2) whether maternal antibodies can protect bank vole offspring against *B. afzelii*-infected *I. ricinus* ticks, and (3) whether this maternal antibody protection is specific for the strain of *B. afzelii*.

## Materials and Methods

### Bank voles, *Ixodes ricinus* ticks and *Borrelia afzelii*

In 2014, we used field-captured bank voles to establish a breeding colony at the University of Neuchâtel [30]. The bank voles used in this study were from the third and fourth lab-born generation and are therefore free from tick-borne pathogens. The *I. ricinus* ticks came from a laboratory colony established in 1978 at the University of Neuchâtel. During the study, the bank voles were maintained in individual cages and were given food and water *ad libitum*. Bank voles were experimentally infected via tick bite with one of two isolates of *B. afzelii*: NE4049 and Fin-Jyv-A3. These two isolates carry two different *ospC* alleles, A10 and A3, which code for two different variants of outer surface protein C (OspC), which is an immunodominant antigen. We have previously shown that immunization with recombinant OspC A10 and A3 induces strain-specific protective antibody responses in laboratory mice [21]. NE4049 was isolated from an *I. ricinus* tick in Switzerland, has multi-locus sequence type (MLST) 679, and strain ID number 1887 in the *Borrelia* MLST database. Fin-Jyv-A3 was isolated from a bank vole in Finland, has MSLT 676, and strain ID number 1961 in the *Borrelia* MLST database. Our previous work has shown that these two strains are highly infectious to both rodents and *I. ricinus* ticks [21,31,32,33].

### Ethics statement and animal experimentation permits

This study followed the Swiss legislation on animal experimentation. The commission that is part of the ‘Service de la Consommation et des Affaires Vétérinaires (SCAV)’ of the canton of Vaud, Switzerland evaluated and approved the ethics of this study. The SCAV of the canton of Neuchâtel, Switzerland issued the animal experimentation permits for the study (NE02-2018) and for the maintenance of the *I. ricinus* tick colony on vertebrate hosts at the University of Neuchâtel (NE05-2014).

### Creation of *I. ricinus* nymphs infected with *B. afzelii*

The nymphs used for the experimental infections were created as follows. BALB/c mice were infected with *B. afzelii* strain NE4049 or Fin-Jyv-A3 via tick bite. At 4 weeks post-infection (PI), the mice were infested with larval ticks from our *I. ricinus* colony. The engorged larval ticks were stored in individual tubes and allowed to molt into nymphs. A random sample of nymphs was tested to determine the infection prevalence, which was 77.9% (67 infected nymphs/ 86 total nymphs) for NE4049 and 91.8% (67 infected nymphs/ 73 total nymphs) for Fin-Jyv-A3. Larval *I. ricinus* ticks were also fed on uninfected BALB/c mice to create uninfected control nymphs.

### Infectious challenge of the bank vole mothers

Five-week-old female bank voles were randomly assigned to one of two experimental groups: control (n = 9) and infected with *B. afzelii* strain NE4049 (n = 11). Each female in the control group was infested with 4 uninfected nymphs; each female in the infected group was infested with 4 nymphs infected with strain NE4049. At 5 weeks PI, a blood sample and an ear tissue biopsy were taken from each female to confirm their infection status. Females were coupled with different males at 2 and 6 weeks PI, and offspring from the first successful coupling was used in the present study. Seven control mothers and 6 *B. afzelii*-infected mothers produced a total of 22 offspring and 20 offspring, respectively (Table S1). At 18 weeks PI, the mothers were sacrificed using CO_2_ asphyxiation and the following organs were aseptically dissected: bladder, left ear, right ear, left rear joint, and right rear joint. The tissue samples were stored at –80°C until further analysis.

### Rearing the bank vole offspring

At 21 days post-birth (PB), offspring were separated from their mothers and moved to individual cages. At 34 days PB, a blood sample and an ear tissue biopsy were taken from each of the 42 offspring. The blood samples were tested for maternal IgG antibodies against *B. afzelii*. The ear tissue biopsies were tested to confirm that there was no mother-to-offspring transmission of *B. afzelii*. As the offspring from the uninfected mothers and the infected mothers are expected to test negative and positive for maternal antibodies (MatAb), they will hereafter be referred to as the MatAb- and MatAb+ offspring, respectively.

At 35 days PB, the offspring were challenged with *I. ricinus* nymphs that were infected with either strain NE4049 or strain Fin-Jyv-A3 (see below).

### Infectious challenge of the bank vole offspring

To test whether maternal antibodies provide strain-specific protection, the MatAb-offspring (n = 22) and the MatAb+ offspring (n = 20) were challenged via tick bite with strain NE4049 or strain Fin-Jyv-A3. Offspring were assigned to balance sample sizes and family effects among the four combinations of MatAb status and challenge strain, which were as follows: MatAb-/NE4049 (n = 9), MatAb-/Fin-Jyv-A3 (n = 11), MatAb+/NE4049 (n = 10), and MatAb+/Fin-Jyv-A3 (n = 10). The remaining 2 MatAb-offspring were challenged with uninfected nymphs as controls. The infectious tick bite challenge for the offspring was the same as for the mothers. Five-week-old offspring were challenged with 4 nymphs infected with either strain NE4049 or strain Fin-Jyv-A3. The engorged nymphs were collected and tested for *B. afzelii* to confirm that each offspring had been infested with at least one infected nymph (Tables S3 and S4). At 5 weeks PI, a second blood sample and a second ear tissue biopsy were taken from each of the 42 offspring to confirm their infection status. At 10 weeks PI, the offspring were sacrificed using CO_2_ asphyxiation and the following organs were aseptically dissected: bladder, left ear, right ear, left rear joint, right rear joint, ventral skin, and dorsal skin. Tissue samples (20–25 mg) from the bladder, left ear, and left rear joint were tested for the presence of *B. afzelii* using qPCR (see below). Tissue samples from the right ear, right rear joint, ventral skin, and dorsal skin were cultured in BSK-II medium (see below).

### Infection status of the bank voles

A bank vole was considered as having been successfully challenged with *B. afzelii* if at least one engorged *B. afzelii*-infected nymph was collected and/or if it developed a systemic infection following the infectious tick challenge. A bank vole was defined as having a systemic infection with *B. afzelii* if it tested positive for more than one of seven criteria: (1) presence of *B. afzelii*-specific IgG antibodies, (2) presence of OspC-specific antibodies, detection of *B. afzelii* spirochetes in (3) ear biopsy at 35 days PI, (4) bladder at 70 days PI, (5) left ear at 70 days PI, (6) left rear joint at 70 days PI, and (7) culture of live spirochetes from dissected organs at 70 days PI.

### *Borrelia*-specific qPCR and *ospC*-specific qPCR

The *B. afzelii* infection status of the engorged nymphs and the bank vole tissue samples was tested using qPCR. The DNA was extracted from the engorged nymphs and the bank vole tissue samples as previously described [30]. The qPCR assay targets a 132 bp fragment of the *flagellin* gene of *B. burgdorferi* sl and was performed as previously described [30]. The strain identity of the engorged nymphs and the offspring ear biopsies was confirmed using a strain-specific qPCR [33]. This qPCR targets a 143 bp fragment of the *ospC* gene and uses two different probes that detect either *ospC* allele A3 or *ospC* allele A10 and was performed as previously described [33].

### *Borrelia*-specific ELISA and OspC-specific ELISA

The serum samples of the voles were tested for the presence of *B. afzelii*-specific IgG antibodies using a commercial ELISA assay as previously described [30]. The maternally transmitted OspC-specific IgG antibody response in the offspring before the infectious challenge (34 days PB) was measured using a homemade ELISA with recombinant OspC (rOspC) proteins A3 and A10 [21]. 96-well tissue culture plates were coated overnight at 4°C with rOspC proteins A3 and A10 (1 μg of protein per well). Wells were washed three times with PBS-Tween 0.1% between each step. The plate was incubated with a BSA 2% blocking solution for 2 hours, followed by the bank vole serum samples (diluted 1:100 in 1x PBS) for 45 minutes, and the secondary antibody for 45 minutes (diluted 1:5000 in 1x PBS). The secondary antibody was a goat anti-*Mus musculus* IgG conjugated to horseradish peroxidase. After adding 100 μl of TMB, we measured the absorbance at 652 nm every 2 minutes for one hour using a plate reader (Synergy HT, Multi-detection plate reader, Bio-Tek, United States). The strength of the IgG antibody response against the rOspC antigens was determined by integrating the area under the absorbance versus time curve.

### Culture of *B. afzelii* spirochetes from bank vole tissues

To demonstrate that the bank vole offspring were infected with live *B. afzelii*, tissue biopsies were cultured in BSK-II media. Tissue biopsies from the skin (ventral skin and/or dorsal skin), right ear, and right rear joint were placed in individual tubes for each of the 42 offspring. The culture tubes were kept in an incubator at 34°C and were screened for live spirochetes over a period of 4 weeks using a dark field microscope.

### Statistical analysis

All statistical analyses were done in R version 1.0.143 (R Development Core Team 2015-08-14). The IgG antibody response is measured in absorbance units and was log10-transformed to improve the normality of the residuals. All means are reported with their 95% confidence intervals (95% CI).

### Maternal infection status and maternal antibody transmission

To test whether the mother bank voles developed an IgG antibody response against *B. afzelii* at 5 weeks PI, we compared this variable (log10-transformed) between infected mothers and uninfected mothers using an independent two samples t-test. To test whether the pre-infection blood sample (at 34 days PB) of the offspring contained maternally transmitted *B. afzelii*-specific IgG antibodies, we compared this variable (log10-transformed) between the MatAb-offspring and the MatAb+ offspring using an independent two samples t-test.

The specificity of the maternal antibodies in the pre-infection blood sample (at 34 days PB) of the offspring was measured as the strength of the IgG antibody response against OspC antigens A3 and A10. We calculated an OspC A10 specificity ratio for each offspring by dividing the IgG antibody response against rOspC A10 by the IgG antibody response against rOspC A3. We compared the log10-transformed OspC A10 specificity ratio between the MatAb+ offspring and the MatAb-offspring using an independent two samples t-test.

### Maternal antibody protection and strain specificity

We tested whether the maternal antibodies protected offspring against infection with *B. afzelii* and whether this protection was strain-specific. Offspring were classified as being uninfected (0) or infected (1) depending on the 7 infection criteria. Offspring infection status was modeled using generalized linear models (GLMs) with binomial errors. The two explanatory factors included offspring maternal antibody status (2 levels: MatAb+ and MatAb-) and offspring challenge strain (2 levels: NE4049, Fin-Jyv-A3), and their interaction. Statistical significance of explanatory factors was determined using log-likelihood ratio tests that compared the change in deviance between nested models to a Chi-square distribution.

## Results

### Maternal infection status and maternal antibody transmission

The mean *B. afzelii*-specific IgG antibody response of the infected females (mean = 3811, 95% CI = 2692–5395) was 7.4 times higher than the uninfected females (mean = 512, 95% CI = 371–706), and this difference was significant (Figure S1; t = −9.335, df = 11, p < 0.001). This result shows that infected mothers developed a strong IgG antibody response against the *B. afzelii* infection. For the offspring blood sample that was taken prior to the infectious challenge, the mean *B. afzelii*-specific IgG antibody response for the MatAb+ offspring (mean = 815, 95% CI = 731–906) was 1.6 times higher than the MatAb-offspring (mean = 511, 95% CI = 459–566) and this difference was significant (Figure 1; t = −5.589, df = 39, p < 0.001). This result shows that the MatAb+ offspring received maternal antibodies from their *B. afzelii*-infected mothers. The OspC A10 specificity ratio of the maternal IgG antibody response was 3.07 times higher in the MatAb+ offspring than the MatAb-offspring, and this difference was significant (Figure S3; t = −10.015, df = 39, p < 0.001). This result shows that the maternal IgG antibodies in the MatAb+ offspring reacted more strongly with rOspC A10 than rOspC A3 compared to the MatAb1 offspring.

**Figure 1.**
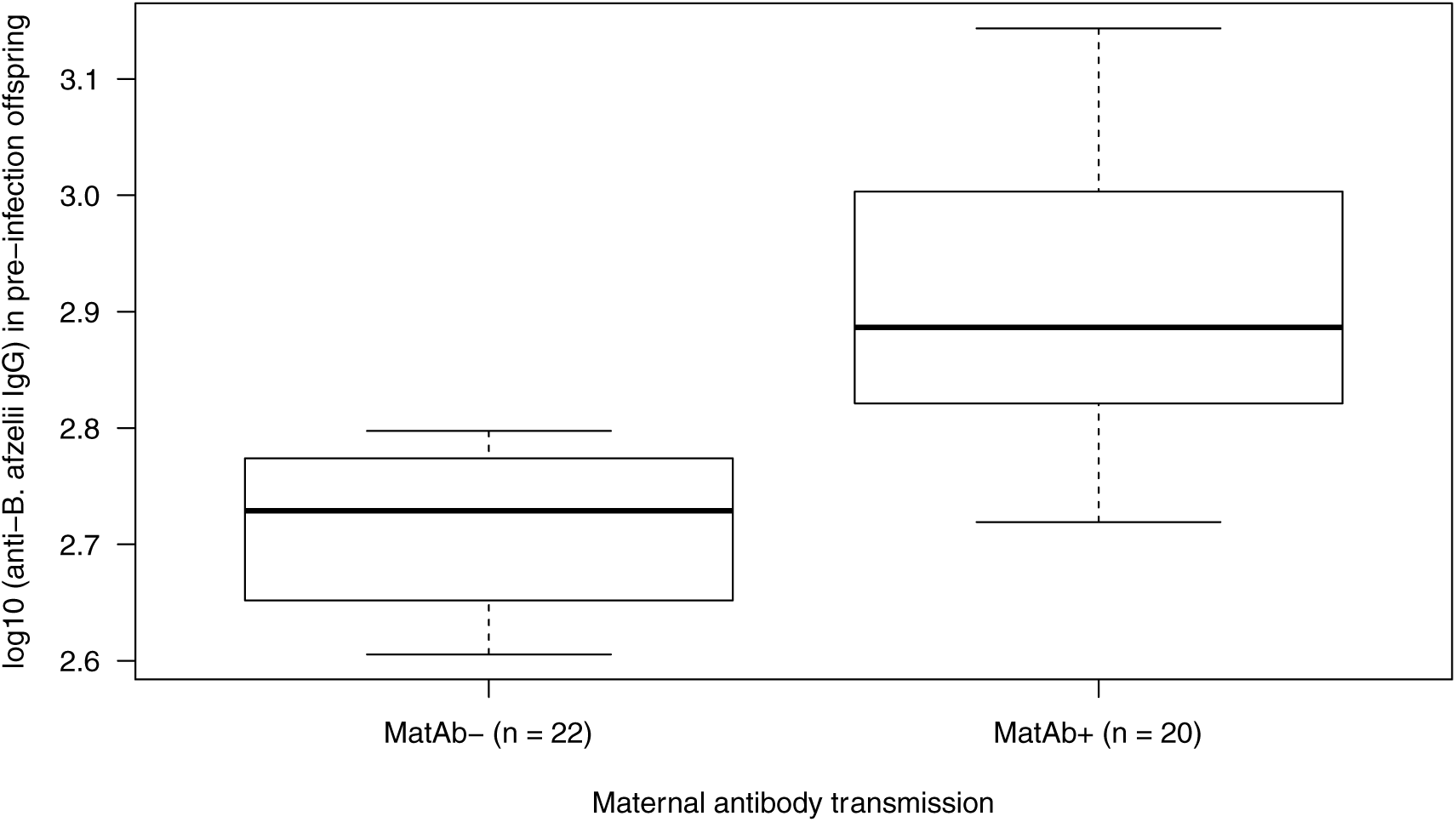
The maternally transmitted *B. afzelii*-specific IgG antibody response was significantly higher in the MatAb+ offspring (n = 20) than the MatAb-offspring (n = 22). The MatAb- and the MatAb+ are the offspring of 7 uninfected control mothers and 6 *B. afzelii*-infected mothers, respectively. The strength of the *B. afzelii*-specific maternal IgG antibody response was measured using a commercial Lyme borreliosis ELISA. Shown are the medians (black line), the 25th and 75th percentiles (edges of the box), the minimum and maximum values (whiskers), and the outliers (circles).

**Figure 2.**
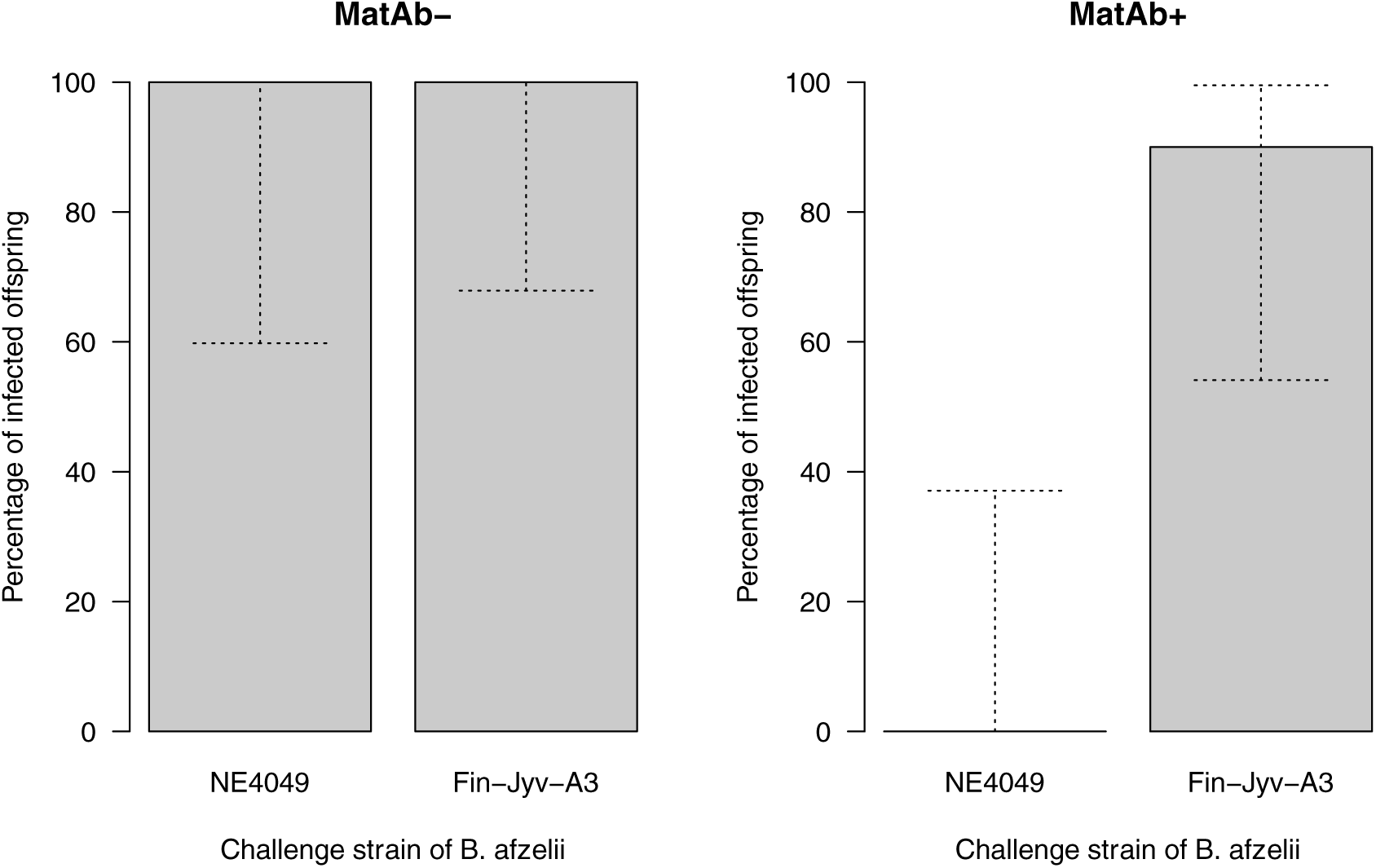
The percentage of infected offspring depends on the maternal antibody status and the challenge strain. The MatAb-(left panel) and MatAb+ (right panel) refer to the offspring from the uninfected control mothers and the mothers infected with *B. afzelii* strain NE4049, respectively. The offspring were challenged via tick bite with either *B. afzelii* strain NE4049 or *B. afzelii* strain Fin-Jyv-A3. The MatAb-offspring were equally susceptible to both strains. The MatAb+ offspring were protected against the maternal strain (NE4049) but not the new strain (Fin-Jyv-A3). The grey solid bars show the means and the stippled bars show the 95% confidence intervals.

### Offspring infection status following the infectious challenge

Before the infectious challenge (34 days PB), the ear tissue biopsies of all offspring tested negative for *B. afzelii* indicating that there was no mother-offspring transmission of the pathogen. The infectious challenge was successful: we collected at least one *B. afzelii*-infected nymph from 38 of the 40 offspring that were challenged with infected ticks. The 2 offspring for which no infected ticks were collected were excluded from the analysis (Tables S3 and S4). The final sample sizes were therefore 8, 11, 9, and 10 offspring for groups MatAb-/NE4049, MatAb-/Fin-Jyv-A3, MatAb+/NE4049, and MatAb+/Fin-Jyv-A3, respectively. The infection status of the offspring was unambiguous; infected offspring tested positive for at least 5 of the 7 infection criteria. In contrast, almost all of the uninfected offspring tested negative for all of the 7 infection criteria; one individual tested positive for 1 infection criterion (Tables S3 and S4).

The analysis of offspring infection status found a highly significant interaction between maternal antibody status and challenge strain (GLM: Δ df = 1, Δ χ^2^ = 71.659, p < 0.001). All of the MatAb-offspring became infected regardless of whether they were challenged with strain NE4049 (100.0% = 8 infected /8 total) or strain Fin-Jyv-A3 (100.0% = 11 infected /11 total). This result shows that both strains were highly infectious to naive offspring. The MatAb+ offspring were perfectly protected against strain NE4049 (0.0% = 0 infected /9 total), but almost completely susceptible to strain Fin-Jyv-A3 (90.0% = 9 infected /10 total), and this difference was significant (χ^2^ = 11.992, df = 1, p < 0.001). This result shows that maternal antibodies only protected offspring against the strain with which the mother had been infected.

## Discussion

### Maternal antibodies are protective and strain-specific

Our study provides the first experimental evidence that maternally transmitted antibodies protect offspring against infection with *B. burgdorferi* sl in an important reservoir host. Earlier studies on a marine Lyme borreliosis system that consists of *B. garinii* and sea birds had shown a positive correlation in antibody concentrations between mothers and their chicks [24]. However, in this sea bird system there was no proof that the maternally transmitted antibodies actually protected the chicks against infection with *B. garinii*. Our study also shows that the protection afforded by the maternal antibodies is highly strain-specific. Offspring from infected mothers were 100% protected against the *B. afzelii* strain to which their mothers had been exposed, but they were highly susceptible to a *B. afzelii* strain to which their mothers had not been exposed. Numerous studies have shown that local populations of *B. burgdorferi* sl contain community of strains that circulate in the same reservoir host and tick populations [12,13,14,15,16]. Theoretical models have shown that strain-specific antibody responses are important for structuring pathogen populations into communities of antigenically distinct strains [6,7,8,9]. Numerous Lyme disease researchers have suggested that the host immune response against the immunodominant OspC antigen could drive the population structure of *B. burgdorferi* sl pathogens [11,12,14,34,35]. The results from our study suggest that the trans-generational transfer of antibodies in vertebrate reservoir hosts could play a critical role in controlling the epidemiology of multi-strain vector-borne pathogens.

### Duration of protection of maternal antibodies

Newborn rodents can take several weeks to develop active immunity [4]. During this period, maternally transmitted antibodies can protect the offspring for 6 to 10 weeks [4,36]. In the present study, we showed that maternally transmitted antibodies protected bank vole offspring at 5 weeks post-birth. A previous study on bank voles found that maternally transmitted antibodies against Puumala hantavirus can protect offspring for a period of two and a half months post-birth [36]. Numerous studies on wild rodents (including bank voles) have shown that sub-adults have a lower prevalence of infection with *B. burgdorferi* sl than adults [19,20,29,37]. The common explanation is that adults have had more time than sub-adults to be exposed to infected ticks. Our study suggests that maternal antibodies may also help to reduce the prevalence of *B. burgdorferi* sl in sub-adult rodents.

### Importance of maternal antibodies for the ecology of Lyme borreliosis

Previous studies on wild bank vole populations in Finland have shown that maternally transmitted antibodies are important for the epidemiology of the Puumala Hantavirus [5]. In contrast to our study, these studies did not investigate strain-specific antibody responses because there is limited antigenic variation in the Puumala Hantavirus [5]. The ecology of Lyme borreliosis suggests that maternally transmitted antibodies could be important for controlling the epidemiology of *B. burgdorferi* sl pathogens in nature. The search for a blood meal by the tick vector, the resultant transmission of *B. burgdorferi* sl, and the reproduction of the rodent host all occur at the same time of the year [11,17]. *Ixodes* nymphs, which transmit *B. burgdorferi* sl, search for reservoir hosts from the spring to the autumn [38,39]. Over the course of the transmission season, the reservoir host population builds up acquired immunity to *B. burgdorferi* sl [19,20]. For example, a study on a population of white-footed mice (*Peromyscus leucopus*) in Connecticut found that 93% of all mice were seropositive for *B. burgdorferi* by the end of August [20]. This observation suggests that at the end of the summer, the majority female rodents are transferring protective antibodies to their offspring. The phenology of nymph questing and *B. burgdorferi* sl transmission coincides with rodent reproduction. For example, the bank vole breeding season is from the spring until early autumn with seasonal fluctuations [17]. In summary, the seasonal buildup of acquired immunity in mothers suggests that there would be high transmission of maternal antibodies to offspring, which would protect offspring from infection with *B. burgdorferi* sl (see below).

### OspC and the mechanism of strain-specific immunity

Our study also showed that the protection of the maternal antibodies was highly strain-specific and most likely mediated by the OspC antigen. The *ospC* is the most polymorphic gene in the genome of *B. burgdorferi* sl [12,13,14] and encodes for outer surface protein C (OspC). OspC is critical for the establishment of infection in the vertebrate host [40,41,42]. Studies have shown that OspC induces a strain-specific antibody response that protects rodents from tick bite [21,22,23,43]. The two *ospC* alleles used in this study (A3 and A10) have a genetic distance of 23.19% and an amino acid distance of 62.57%. We had previously shown in a vaccination trial that rOspC proteins A3 and A10 induce strain-specific protection against strains of *B. afzelii* carrying the corresponding *ospC* alleles [21]. In the present study, we showed that maternal infection with *B. afzelii* strain NE4049, which carries *ospC* allele A10, resulted in the presence of IgG antibodies in the offspring that reacted much more strongly with rOspC A10 than rOspC A3. Taken together, our results suggest that maternal antibodies are highly strain-specific and that OspC is a critical antigen for this specificity.

### Importance of maternal antibodies for population structure of *ospC* type strains

Long-term field studies on *B. afzelii* in tick populations and rodent populations have shown that the community of strains carrying different *ospC* major groups (oMGs) was stable over more than a decade, with some strains an order of magnitude more common than others [15,16]. Strains that were common in the field had higher rates of host-to-tick transmission in laboratory studies [15,44]. An important question is why these high transmission strains do not eliminate the low transmission strains. The *ospC* polymorphism is maintained by balancing selection and two alternative hypotheses, multiple niche polymorphism (MNP) and negative frequency-dependent selection (NFDS), have been proposed [12,13,35]. Under MNP, the different oMG genotypes are adapted to different host species and the frequency of each oMG genotype depends on the abundance of their respective host species [13,35]. Under NFDS, the immune system of the vertebrate host is responsible for controlling the frequencies of the different oMG types. This model suggests that the host immune system is more efficient at targeting the more common oMG strains, and that the rare oMG strains therefore have a selective advantage [12,14,45]. The present study suggests that balancing selection could result from the maternal transfer of OspC-specific antibodies. Under this mechanism, acquired immunity in the mothers would build up faster against the common oMG strains than the rare oMG strains, and the offspring would be more likely to have protective maternal antibodies against the former than the latter. The seasonal trans-generational transmission of strain-specific acquired immunity could prevent the common oMG strains from eliminating the rare oMG strains.

## Conclusions

We used experimental infections with a common Lyme disease pathogen (*B. afzelii*) and its natural reservoir host (the bank vole) to show that females transmit maternal antibodies to their offspring. These maternal antibodies were completely protective against the strain that the mother had encountered, but provided no protection against a different strain. The immunodominant OspC antigen appears to mediate this strain-specific maternal antibody response. The inter-generational transfer of protective strain-specific antibodies could have important implications for the epidemiology of multiple strain pathogen populations in the field.

Future studies should investigate whether maternal antibodies are important for protecting other important reservoir host species against *B. burgdorferi* sl, such as the white-footed mouse (*Peromyscus leucopus*) in North America. They should investigate the duration of protection, the mechanism of antibody transfer (e.g. via the placenta or via milk), and whether females infected with multiple strains transmit antibody responses that are protective against each of those strains. Additional studies are needed to test whether OspC is solely responsible for the strain-specific immunity or whether other *B. burgdorferi* sl antigens are involved. Finally, theoretical models are needed to investigate how the maternal transfer of antibodies in the reservoir host population would influence the epidemiology of this multi-strain tick-borne pathogen.

## Acknowledgements

This work was supported by a grant from the Swiss National Science Foundation to Maarten Voordouw (FN 31003A_141153). We thank Jacob Koella for advice on the experimental design.

## Author contributions

A.G.-C. and M.J.V designed the study. A.G.-C., V.H, A.S and O.R. performed the experimental infections. A.G.-C., A.S and O.R. performed the molecular work. A.G.-C., A.S. and D.G. created the *B. afzelii*-infected nymphs. M.J. created the recombinant proteins. A.G.-C. analysed the data. A.G.-C. and M.J.V wrote the manuscript. All authors read and approved the final version of the manuscript.

Competing interests: The authors declare no competing of interest.

## Supporting Information Legends

The file titled “**Supporting information MatAb_v14.docx**” contains the raw data from this study and nine sections that contain additional results.

